# An Efficient Microfluidic Platform for Organ-Specific Cyclic Multiaxial Loading and Differentiation of Progenitor Cells

**DOI:** 10.64898/2026.11.01.763270

**Authors:** Nithya Murugesan, Silpa Arkat, Doss Aristotle, Supratim Saha, Arnov Paul, Soma Guhathakurta, Ajit Mullasari, Prashant Shah, Shrishu R Kamath, Nitish R Mahapatra, Sarit K. Das

## Abstract

Mechanical loading is a fundamental regulator of tissue development, maturation, and function, yet existing *in vitro* systems predominantly apply uniaxial load/strain to cell-laden hydrogels, limiting their ability to reproduce physiologically relevant multiaxial on-chip tissue mechanics. Here, we present an engineered system to apply controlled multiaxial mechanical loading to real scaffolds with undifferentiated cells. The system enables programmable cyclic stretching and relaxation at the cell scaffold level and is compatible with both cell-laden native scaffold and engineered soft tissues. We have quantified the strain field experienced by the tissue with distinct mechanical loading periods using computation simulations and validated these with systematic experiments, demonstrating reproducible and various organ-relevant multiaxial strain profiles. As proof of concept, we applied the approach to human cardiac tissues, where cyclic multiaxial mechanical loading of mononuclear cells laden native tissue scaffold promoted cardiac structural organization and supports the emergence of spontaneous contractile activity. This approach provides a generalizable method for reproducing physiologically relevant tissue mechanics *in vitro* and enables systematic investigation of mechano-regulation across a plethora of human tissues.

## INTRODUCTION

The evolving field of biomaterials and manipulative effort by microfluidic field investigates incorporating precise control over cellular micro environmental factors, such as microarchitecture, in engineered tissues with the ultimate goal of directing physiological tissue function and maturation^1,2^. *In vivo*, most soft tissues experience cyclic stretching and relaxation in more than one direction, giving rise to complex, multiaxial strain fields that influence cell behaviour and extracellular matrix reorganization^3^. Reproducing these mechanical cues *in vitro* is therefore essential for studying mechano-regulation and for generating functionally relevant human tissue models that can be utilized for drug testing and other therapeutic and diagnostic purposes at the tissue as well as cellular levels.

Current *in vitro* mechanical stimulation in microfluidic platforms have provided valuable insights but remain limited in their ability to reproduce physiological dynamic tissue mechanics^4–7^. Most reported platforms apply uniaxial/multiaxial strain, often to cell-laden hydrogels, which simplifies actuation and control but does not capture the multiaxial loading (longitudinal, circumferential, and radial strains), of organ specific nature experienced by native tissues such as myocardium, lung, blood vessels, intestine, etc.^8–13^. In addition, hydrogel-based approaches often lack native tissue architecture and mechanical heterogeneity, constraining their relevance in comparison to realistic normal tissue models^12,14,15^.

Several approaches have attempted to increase the mechanical complexity by combining uniaxial/multiaxial stretch with patterned substrates or anisotropic scaffolds^16^. However, these strategies still impose directionally constrained loading and do not allow independent control or validation of multiaxial strain at the native tissue scale. As a result, there remains a need for a general method that can apply controlled reproducible microenvironment for achieving the proposed goal of functional engineered tissue.

Here, we propose an innovative method in which acellular native scaffold is wrapped around an inflatable balloon structure integrated within a microfluidic system. Cyclic inflation and deflation of the balloon generate controlled dynamic stretching and relaxation of the surrounding tissue, producing multiaxial mechanical loading that more closely resembles physiological deformation. This method enables tissue-level actuation in the microfluidic platform without the requirement of hydrogel encapsulation methods^15^ and is compatible with a range of relevant human tissues exhibiting diverse stretching-relaxation cycle frequencies.

We quantify the strain fields generated within the wrapped tissues using the computational simulations and validate these simulations though experimental measurements, demonstrating reproducible and tunable multiaxial strain across multiple loading regimes. As a proof of concept, we apply this method to human acellular myocardial scaffold (AMS) laden with mononuclear cells (progenitors), where cyclic multiaxial mechanical loading supports myocardial structural organization and are able to bring out a break through with the emergence of spontaneous contractility of the mononuclear cells differentiated cardiac cells. Together, this approach establishes a generalizable method for reproducing physiologically relevant cyclic multiaxial tissue mechanics *in vitro* which not only provides a foundation for systematic investigation of mechanical regulation across human tissues but also effects the differentiation of primary cells with functional transformation.

### Device conceptualization and design rationale

The microfluidic platform was conceptualized to recreate key biomechanical and physiological features of native tissue within an *in vitro* system (Fig. 1a). The design integrates three essential elements: (i) an engineered native scaffold providing a three-dimensional extracellular matrix (ECM), (ii) cyclic multiaxial mechanical stimulation delivered via a balloon catheter to mimic organ specific rhythmic deformation, and (iii) continuous perfusion to sustain cell-laden scaffold viability and metabolic exchange (Fig. 1b). Decellularized cardiac tissue was selected as the scaffold material to preserve native ECM architecture, and bioactive cues critical for cell differentiation, maturation in targeted nutrient media. Mechanical stimulation was achieved using a centrally positioned balloon catheter capable of programmable inflation and deflation, enabling controlled cyclic stretch and relaxation analogous to the physiological conditions. Continuous perfusion through adjacent microchannels supported nutrient delivery, oxygen transport, and metabolic waste removal, thereby approximating physiological blood perfusion (appropriate for tissue culture system).

The device architecture comprises a central tissue chamber hosting tissue wrapped around balloon catheter flanked by perfusion channels, with optical access for real-time monitoring. Polydimethylsiloxane (PDMS) was selected as the primary material for the device due to its biocompatibility, flexibility, and optical transparency^17^. The catheter balloon, made of medical-grade silicone, was chosen for its elasticity and durability, ensuring reliable performance under repeated actuation over days. The design integrates mechanical, material, and biological principles, providing an innovative and versatile platform for studying cell-ECM behaviour under dynamic conditions. This integrated design enables simultaneous application of cyclic multiaxial mechanical (C-MAM) conditioning and perfusion in a compact, reusable microfluidic platform.

**Figure. 1.**
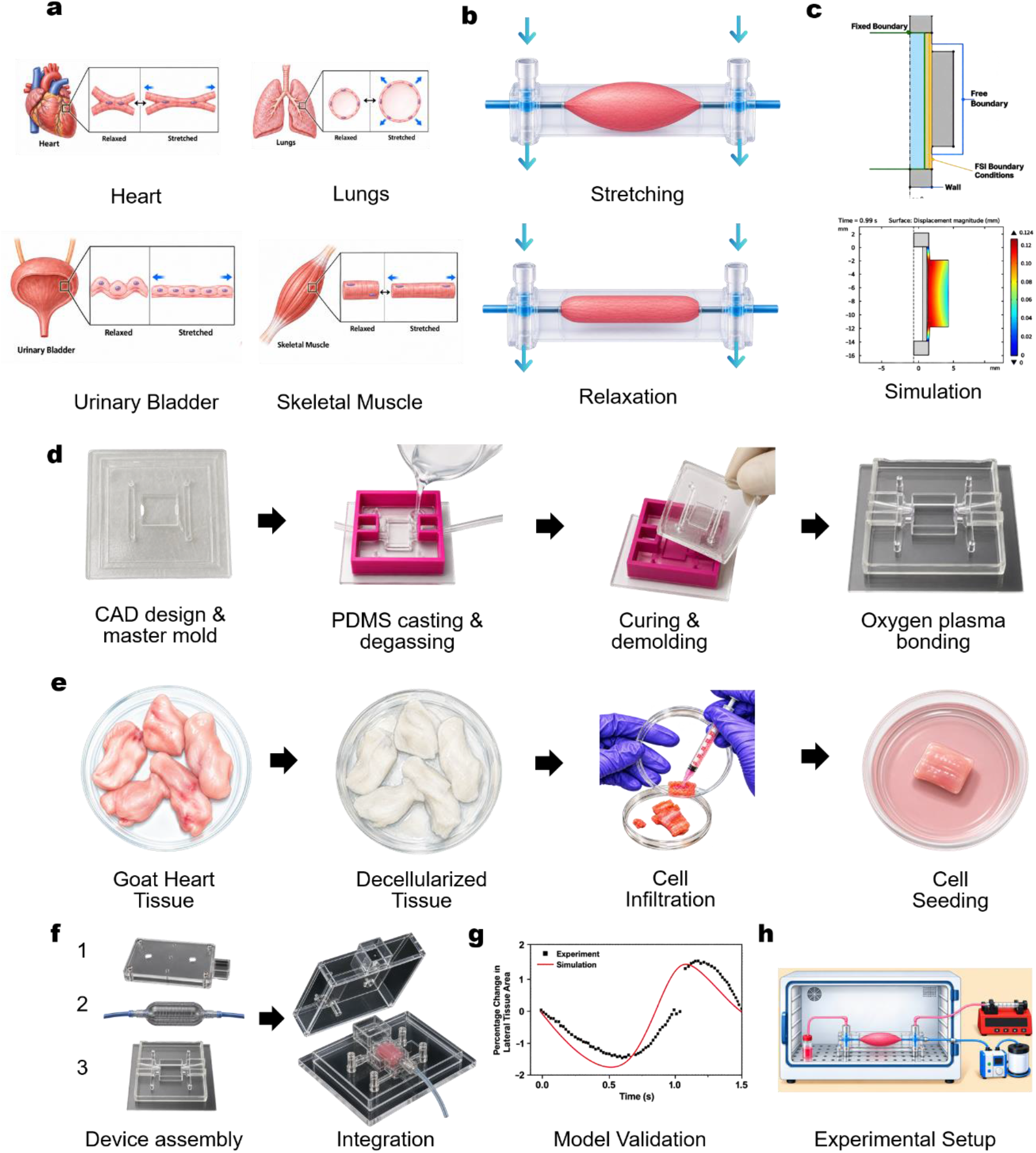
Integrated experimental and computational framework for cyclic mechanical stimulation and tissue engineering of human-organ-inspired tissue constructs. **(a)** Physiological examples of cyclic stretching and relaxation of the heart, lungs, urinary bladder, and smooth muscle tissues and their associated functions. **(b)** Working principle of the microfluidic device, illustrating cyclic inflation and deflation of the tissue construct with bidirectional media perfusion through the lateral microchannels. **(c)** Finite element simulation of tissue deformation under pressure-driven inflation, including the fluid–structure interaction (FSI) boundaries and tissue displacement. **(d)** Device fabrication workflow involving CAD design, high-resolution 3D printing, master mold and cap fabrication, PDMS casting, vacuum degassing, thermal curing, demoulding, oxygen-plasma activation, and PDMS–glass bonding. **(e)** Decellularization of native tissue using sodium deoxycholate treatment followed by repeated washing to obtain an extracellular matrix (ECM) scaffold, cell seeding and infiltration of mononuclear cells into the decellularized ECM scaffold. **(f)** Assembly of the PDMS microfluidic device with the removable cover and balloon catheter **(g)** Experimental validation of the numerical model by comparison of the simulated and experimentally measured temporal variation in lateral tissue area. **(h)** Ex vivo tissue-culture setup under controlled environmental conditions and cyclic mechanical stimulation.

### Device characterization and numerical–experimental validation of cyclic strain

The balloon catheter–assisted actuation system enabled reproducible and tunable C-MAM deformation of acellular tissue constructs across a wide range of frequencies and amplitudes relevant to multiple organ systems. By modulating inflation–deflation duration and applied pressure, the platform generated controlled strain profiles corresponding to physiological deformation regimes observed in the heart, lungs, muscles, and bladder (Figs. 2, 3).

To quantitatively characterize tissue deformation and establish predictive control over strain delivery, numerical simulations were performed using a fluid–structure interaction (FSI) framework. A two-dimensional axisymmetric computational model was developed to represent the coupled interaction between the pressurized air domain within the balloon, the hyperelastic silicone balloon wall, and the surrounding acellular tissue construct (Fig. 2a). Cyclic pressure loading at the balloon inlet resulted in expansion and contraction of the balloon wall, which was transmitted directly to the tissue through a perfectly bonded interface, thereby mimicking experimentally imposed stretching–relaxation. The simulations revealed spatially and temporally resolved strain distributions around the scaffold that closely followed the programmed pressure waveforms (Videos GIF’s 1, 2, 3 and 4). High-frequency, low-duration actuation produced rapid, pulsatile strain patterns characteristic of cardiac tissue deformation and smooth muscles (Figs. 2b, 3b). Intermediate frequencies generated smooth, rhythmic expansion profiles consistent with pulmonary mechanics, while low-frequency, high-amplitude actuation yielded sustained distension analogous to bladder filling dynamics (Figs. 2g, 3g). Across these regimes, simulated strain magnitudes scaled predictably with applied pressure amplitude and cycle duration, demonstrating robust tunability of the mechanical stimulus

**Figure. 2.**
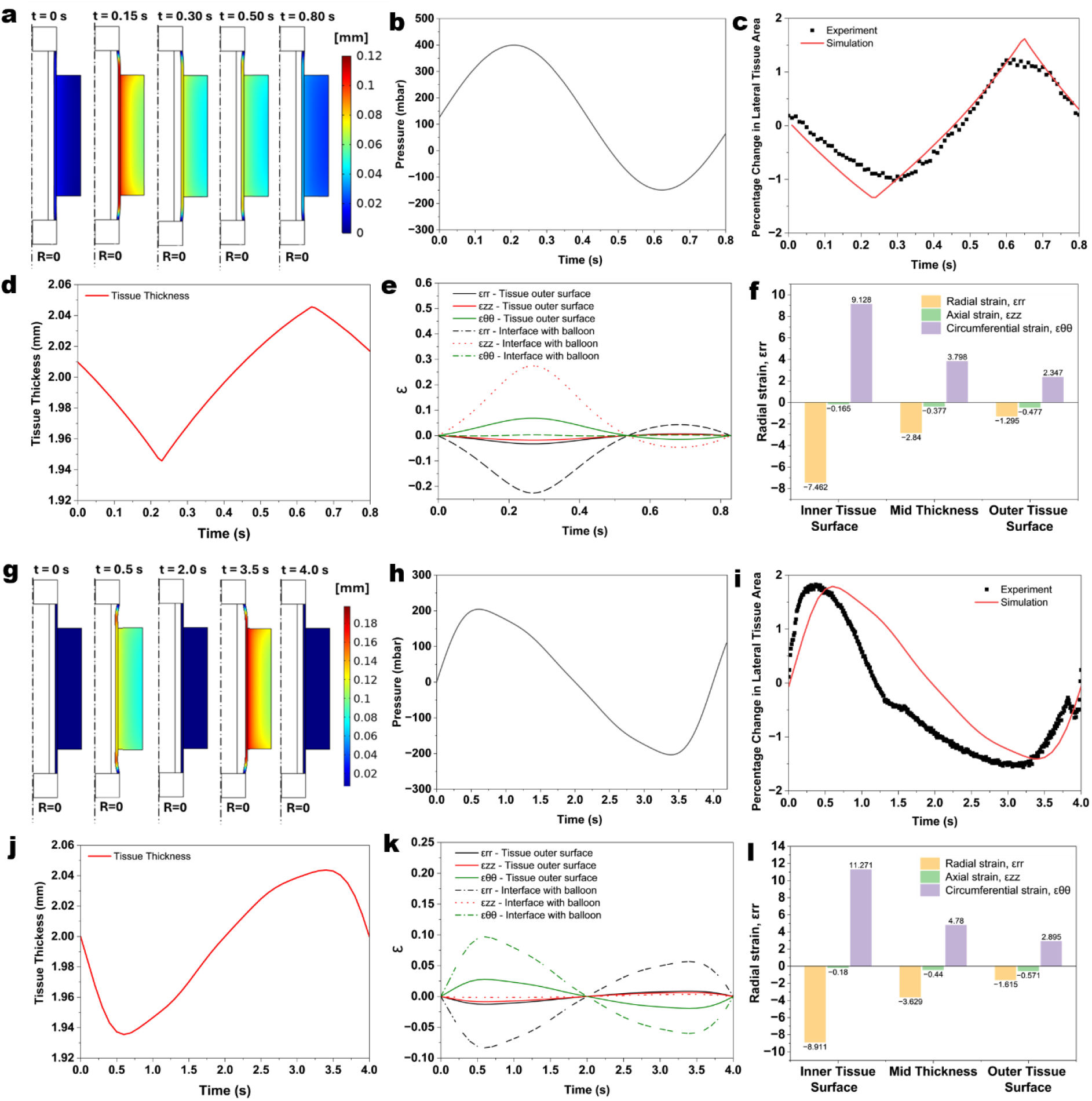
Experimental validation of the numerical simulations and predicted deformation and strain characteristics of heart and lung tissues during cyclic inflation–deflation. **(a–e)** Heart tissue: **(a)** numerical prediction of tissue displacement magnitude and applied pressure during inflation (0.3 s) and deflation (0.5 s); **(b)** comparison of experimentally measured and numerically predicted percentage change in lateral tissue area for model validation; **(c)** numerically predicted temporal variation in tissue thickness; **(d)** temporal evolution of the radial (ε_RR_), axial (ε_ZZ_), and circumferential (ε_ФՓ_) strain components at the tissue outer surface and tissue–balloon interface; and **(e)** through-thickness distribution of the three normal strain components at peak loading ((t=0.2) s). **(f–j)** Lung tissue: **(f)** numerical prediction of tissue displacement magnitude and applied pressure during inflation (2 s) and deflation (2 s); **(g)** comparison of experimentally measured and numerically predicted percentage change in lateral tissue area for model validation; **(h)** numerically predicted temporal variation in tissue thickness; **(i)** temporal evolution of the radial, axial, and circumferential strain components at the tissue outer surface and tissue–balloon interface; and **(j)** through-thickness distribution of the three normal strain components at peak loading ((t=0.6) s).

Experimental validation of the numerical predictions was performed by directly measuring strain experienced by the acellular scaffold during cyclic actuation. A strain sensor connected to a multichannel data acquisition system was affixed to the lateral surface of the tissue scaffold, and strain was recorded under mechanical stimulation designed to replicate organ-specific stretching– relaxation (Fig. S1). The experimentally measured temporal strain profiles closely matched the numerically predicted strain at the corresponding tissue (heart, lungs, smooth muscle and bladder) location (Figs. 2b, 2g, 3b, 3g), confirming accurate transmission of balloon-induced deformation to the scaffold. The experimentally measured lateral-area changes ranged from -2.338 % to 2.082 %, whereas the corresponding numerical predictions ranged from − 1.47022 % to 1.792 %.

Quantitative comparison yielded RMSE values of 0.288, 0.630, 0.5611, and 1.2360 percentage points, with corresponding Pearson correlation coefficients of 0.953, 0.899, 0.8223, and 0.8214, respectively. The good agreement between the experimental and numerical responses established the validity of the computational models resulting in the use of validate numerical model to predict the local deformation and mechanical fields that cannot be directly resolved from the lateral-area measurements. This combined numerical–experimental validation establishes that the platform delivers well-defined, organ-relevant strain profiles. Together, these results demonstrate that the developed microfluidic platform provides physiologically reproducible mechanical stimulation within a single device with precision, qualifying comparative mechanobiology studies across the dynamic organs using acellular scaffolds with targeted cells.

**Figure. 3.**
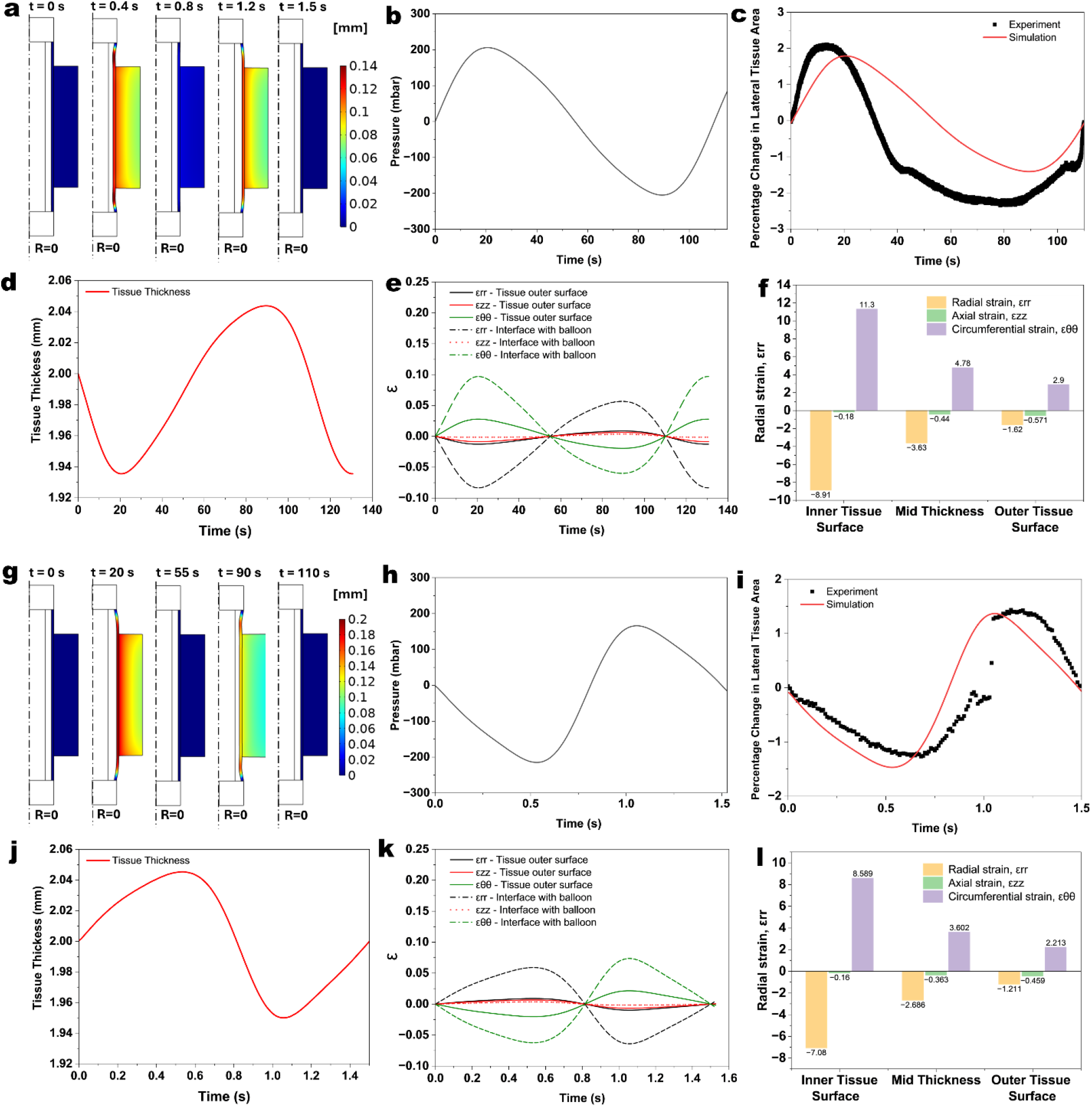
Experimental validation of the numerical simulations and predicted deformation and strain characteristics of smooth-muscle and bladder tissues during cyclic inflation–deflation. **(a–e)** Smooth-muscle tissue: **(a)** numerical prediction of tissue displacement magnitude and applied pressure during inflation (0.5 s) and deflation (1 s); **(b)** comparison of experimentally measured and numerically predicted percentage change in lateral tissue area for model validation; **(c)** numerically predicted temporal variation in tissue thickness; **(d)** temporal evolution of the radial (ε_RR_), axial (ε_ZZ_), and circumferential (ε_ФՓ_)strain components at the tissue outer surface and tissue–balloon interface; and **(e)** through-thickness distribution of the three normal strain components at peak loading ((t=1.05) s). **(f–j)** Bladder tissue: **(f)** numerical prediction of tissue displacement magnitude and applied pressure during inflation (30 s) and deflation (60 s); **(g)** comparison of experimentally measured and numerically predicted percentage change in lateral tissue area for model validation; **(h)** numerically predicted temporal variation in tissue thickness; **(i)** temporal evolution of the radial, axial, and circumferential strain components at the tissue outer surface and tissue– balloon interface; and **(j)** through-thickness distribution of the three normal strain components at peak loading ((t=20) s).

The temporal evolution of the normal strain components demonstrated a multiaxial and spatially heterogeneous deformation response in all four tissue models (Figs. 2d, i and 3d, i). Radial, axial, and circumferential strain components evolved simultaneously during cyclic balloon inflation and deflation, although their relative magnitudes varied according to the tissue type and loading regime. In general, the deformation was more pronounced near the tissue–balloon interface than at the outer tissue surface, indicating that the mechanical interaction with the expanding balloon generated a pronounced through-thickness deformation gradient.

At the peak loading state, the through-thickness strain distributions further demonstrated the spatial heterogeneity of deformation (Figs. 2e, j and 3e, j). In the heart tissue, the radial, axial, and circumferential strains at the tissue–balloon interface were −7.462%, −0.165%, and +9.128%, respectively. At the mid-thickness location, the corresponding values were −2.840%, −0.377%, and +3.798%, while at the outer tissue surface they decreased to −1.295%, −0.477%, and +2.347%, respectively. The corresponding strain distributions for the lung, smooth-muscle, and bladder tissues exhibited the same general through-thickness localization, although the magnitude of the individual strain components varied among the tissue types. Across the four models, the largest radial compression and circumferential extension were generally observed adjacent to the tissue– balloon interface and progressively decreased toward the outer surface.

The numerical stress fields provided additional insight into the local mechanical loading that could not be obtained directly from the experimental deformation measurements. The predicted stresses exhibited pronounced spatial localization near the tissue–balloon interface, where the tissue experienced the greatest deformation. For the heart tissue, the circumferential, radial, and axial stresses at the tissue–balloon interface reached approximately 185 kPa, −13.6 kPa, and 76.4 kPa, respectively. At the mid-thickness location, the corresponding values decreased to approximately 0.380, −0.113, and 0.055 kPa, while at the outer tissue surface they were approximately 0.260, −0.010, and 0.047 kPa, respectively (Table S1). The corresponding stress distributions for the lung, smooth-muscle, and bladder tissues are summarized in Table S2, S3 and S4. Across the four models, the stress magnitude generally decreased from the tissue–balloon interface toward the outer tissue surface, consistent with the observed through-thickness strain gradients.

### Decellularized scaffold validation, cell homing, proliferation, and motion analysis

Decellularization efficiency was assessed by quantifying the residual DNA content (Figs. S1a and b) and by histological analysis using hematoxylin and eosin (H&E), Masson’s trichrome, and Verhoeff–Van Gieson (EVG) staining. Residual DNA content in decellularized scaffolds was reduced by 90% relative to native tissue (Fig. 4a), confirming effective removal of cellular material. H&E staining revealed an absence of nuclei alongside preservation of the overall extracellular matrix (ECM) architecture, when compared with native controls. Masson’s trichrome staining showed loss of cellular components with retention of collagen fibres (blue), while EVG staining confirmed preservation of elastin (black) and collagen (red) within the decellularized cardiac matrix (Fig. S1c). Collectively, these results confirm effective decellularization with retention of key structural ECM components, including collagen I and IV, laminin, and fibronectin.

To evaluate the homing and proliferative capacity of mononuclear cells (MNCs) on the decellularized scaffold, DAPI staining were performed at days 14, 18, 21, and 28 post-seeding. Cell homing peaked at day 21, followed by a decline in proliferative activity at later time points. Notably, MNC proliferation was consistently greater under mechanical stimulation than under static culture conditions. Representative images and their analysis are shown in Figs. 4a and b. FFT analysis of a representative case revealed distinct frequency characteristics between the two conditions as shown in Fig. 4c. Cells exposed to mechanical stimulation exhibited a dominant low-frequency component at approximately 0.1 Hz, indicating slower motion variations over a longer characteristic timescale. In contrast, cells without mechanical stimulation showed a dominant frequency near 1.0 Hz, suggesting faster motion fluctuations. Consistent with this observation, PSD analysis showed predominantly low-frequency power under mechanical stimulation, whereas the unstimulated condition exhibited a broader distribution of spectral components, particularly around 0.8–1.3 Hz and 2.1–2.7 Hz as demonstrated in Fig. 4d. The autocorrelation profiles further demonstrated condition-dependent differences in motion recurrence as illustrated in Fig. 4e. Collectively, this representative case illustrates that mechanical stimulation may shift cellular motion toward slower, longer-timescale and potentially more coordinated dynamics.

**Figure. 4.**
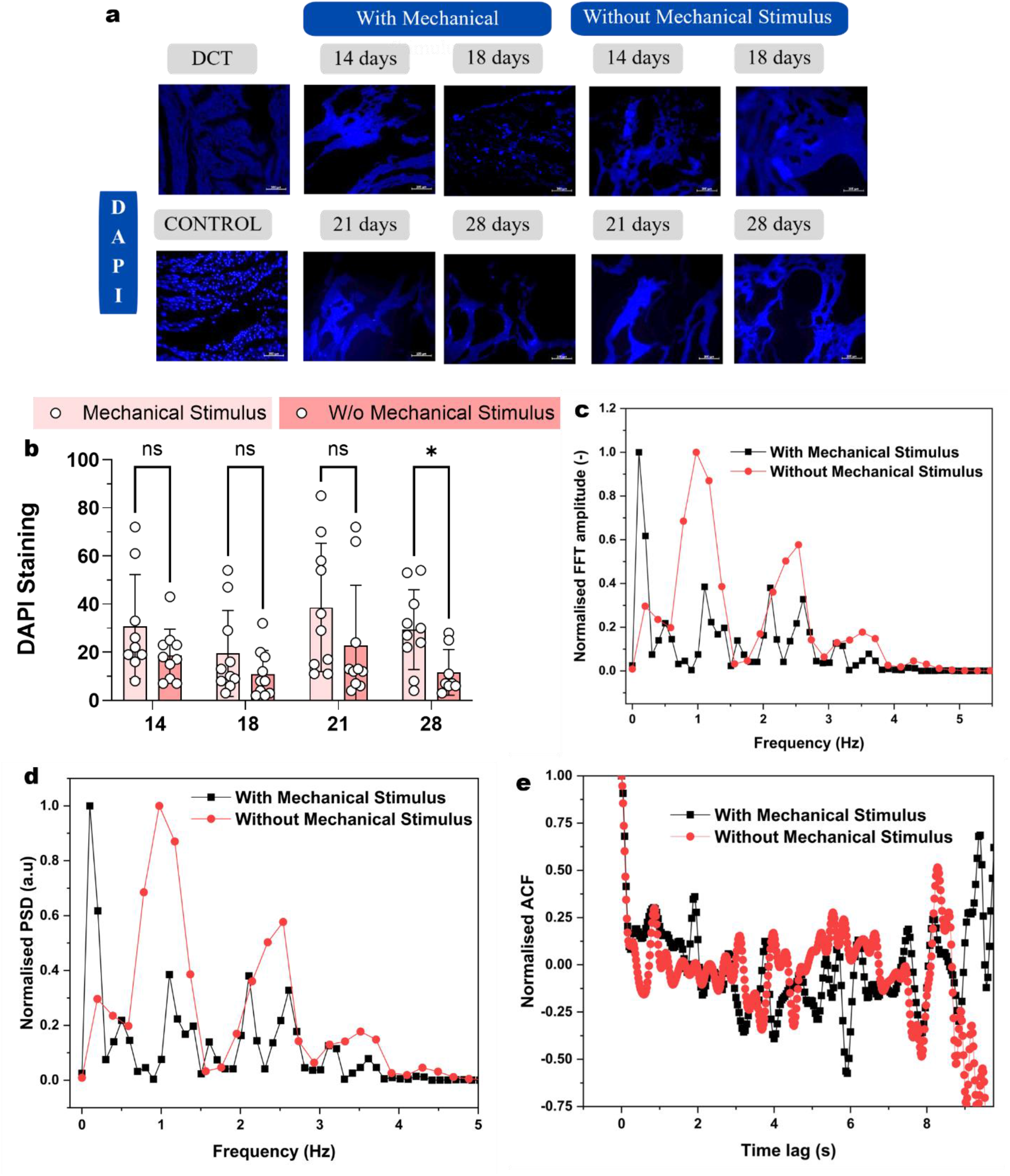
Cell homing, proliferation and motion analysis of differentiated cardiomyocytes under mechanical stimulus. **(a)** Representative DAPI-stained fluorescence images showing nuclei of mononuclear cells (MNCs) homed onto decellularized cardiac scaffolds at 14, 18, 21, and 28 days under static and mechanically stimulated culture conditions. Scale bar, 100 µm. **(b)** Quantification of DAPI-positive nuclei per field of view, expressed as cell number/mm², analyzed using ImageJ software (NIH). For each condition and time point, [n] non-overlapping fields were quantified from [n > 5] independent scaffolds. Data are presented as mean ± s.d. Statistical significance was determined by two-way ANOVA with [Tukey’s/Bonferroni’s] post hoc test for multiple comparisons, considering culture condition (static vs. mechanical stimulation) and time (days 14–28) as independent variables. p < 0.05 was considered statistically significant. **(c)** Fast Fourier transform (FFT) **(d)** Power Spectral density (PSD) **(e)** Autocorrelation coefficient (ACF), of with and without mechanical stimulus.

### Initiation of mechanical stimulation for cardiomyogenic differentiation

Peripheral blood mononuclear cells (PBMCs) were procured commercially (Himedia -HiFiTM Human Peripheral Blood Mononuclear Cells (H-PBMC), Single Donor - 25 million cells /vial - CL003-25), thawed in RPMI medium, and pre-cultured for 12–24 hours. Approximately 2-5 × 10⁶ mononuclear cells (MNCs) were then injected into the processed sterile decellularized scaffolds using 28G insulin needles through multiple micro-punctures, and maintained in low glucose DMEM complete medium for 3 days. The MNC-seeded scaffolds were subsequently transferred into the microfluidic devices by wrapping those around the balloon catheter positioned in the central chamber of microfluidic platform to provide mechanical stimulation. Same set of scaffolds injected with cells without mechanical stimulus were also maintained in separate contraption. Cultures were supplied with fresh conditioned medium on alternate days to ensure a congenial environment for differentiation. Mechanical stimulation (60 per minute) was applied for 2 weeks, after which it was withdrawn, and the cultures were maintained for an additional 2 weeks to promote maturation and observations were made during this period while changing the old media by new nutrient media. The overall workflow is illustrated in the Fig. S2.

### Confirmation of cardiomyogenic differentiation and maturation

Immunofluorescence analysis revealed progressive cardiomyogenic differentiation and maturation of mononuclear cells within the mechanically stimulated scaffolds. Troponin expression, indicative of cardiac contractile apparatus formation, was faint at earlier time points (days 14–18) but became markedly stronger by 21^st^ day (p = 0.05 compared to without stimulus), indicating activation and maturation of cardiac-specific contractile machinery (Fig. 5a). This observation validates the efficiency of the differentiation protocol, reflecting both lineage specification and functional maturation of the cells in MNC-seeded scaffolds. This increase was more pronounced in mechanically stimulated tissues compared to unstimulated controls. Connexin-43 (Cx43) staining revealed temporal maturation of electrical coupling. The immature cardiomyocytes exhibit diffuse or cytoplasmic Cx43 localization, which progressively shifted toward membrane-associated and clustered patterns as the cells mature. In our system, as expected the early-stage cultures (days 14 and 18) exhibited diffuse or cytoplasmic localization, whereas later stages (days 28, p < 0.05), particularly the tissue under mechanical stimulation showed pronounced membrane-associated clustering consistent with functional gap junction formation (Fig. 5b) as cells matured . We also observed progressive increase in the expression of α-sarcomeric actinin (Fig. 5c) at days 21 and 28 (p < 0.05 compared to without stimulus), indicative of enhanced cardiomyogenic differentiation over time. This increase was significantly more pronounced under mechanical stimulation compared with static culture conditions, suggesting that biomechanical cues play a critical role in promoting cytoskeletal and sarcomeric organization within the seeded scaffolds

**Figure. 5.**
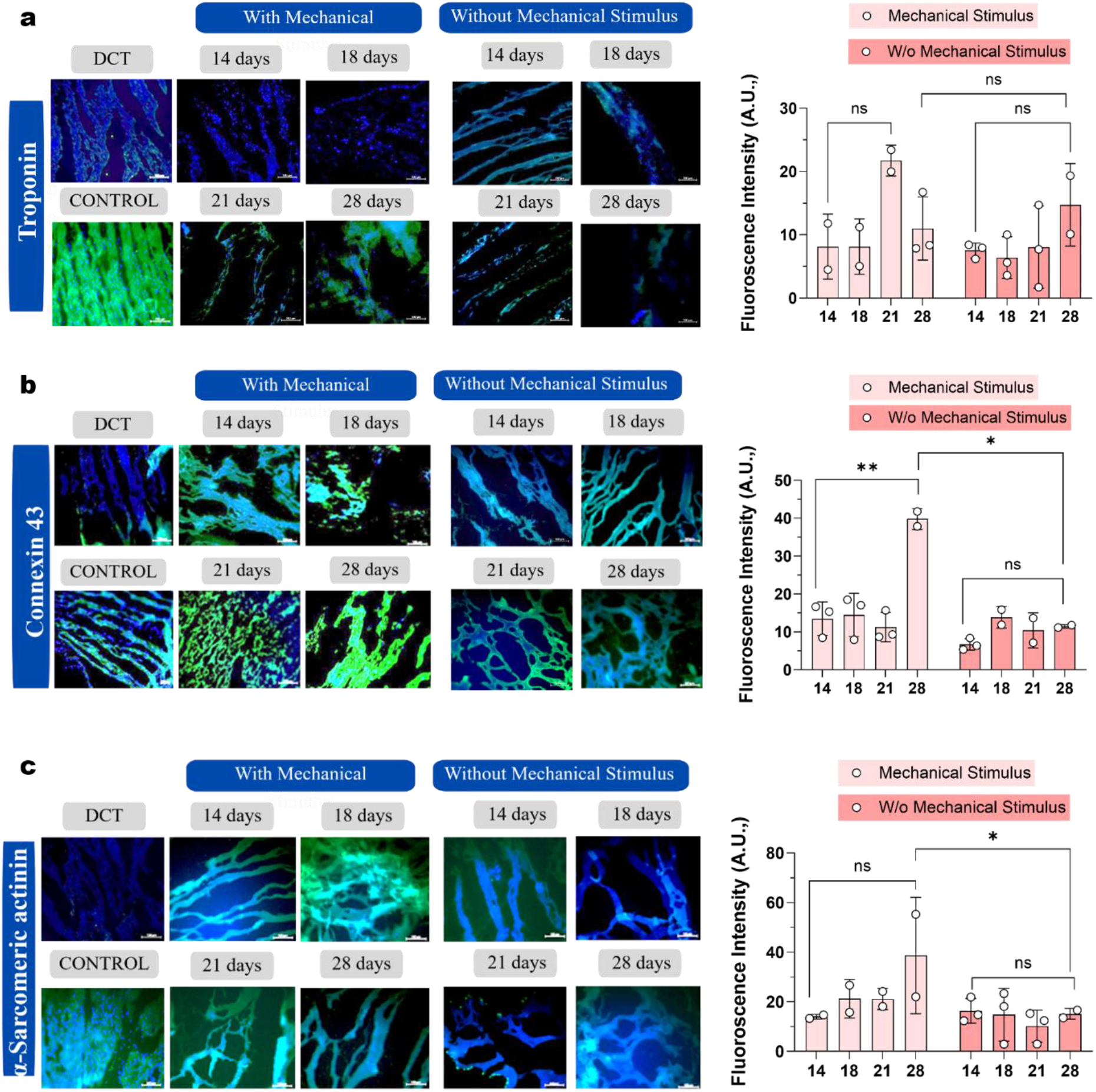
Immunofluorescence analysis of cardiac-specific markers in the presence and absence of mechanical stimulation. **(a-c)** Representative immunofluorescence images and quantification of Troponin, Cx43, alpha-sarcomeric actinin expression across the culture period from day 14 to day 28. Fluorescence intensity for all markers was quantified using ImageJ software (NIH) from [n=3] independent fields per condition and time point across [n=3] biological replicates. Data are presented as mean ± s.d. Statistical significance was assessed by one-way ANOVA with [Tukey’s/Bonferroni’s] post hoc test, considering time and culture condition (static vs. mechanical stimulation) as independent variables (\**P* < 0.05; ns, not significant)

Vimentin expression, characteristic of immature or mesenchymal-like cells, was high at early differentiation stages (day 14) and progressively decreased at later time points (day 18, p = 0.05), consistent with cardiomyocyte maturation (Fig. 6a). The decline in vimentin expression was more evident in mechanically stimulated tissues, supporting the role of cyclic strain in promoting phenotypic maturation. ANP belongs to the so-called “fetal gene program” — a set of genes, including ANP, BNP, and β-myosin heavy chain, that are highly expressed during early cardiac development and are re-activated in adult cardiomyocytes under conditions of mechanical overload, hypoxia, or pathological stress. We observed progressive decline in ANP expression ( p = 0.2 at day 21 compared to without stimulus) that is consistent with its well-established role as a marker of immature or stress-responsive cardiomyocyte phenotypes rather than mature, functionally organized cardiac tissue (Fig. 6b)

To assess the molecular response of beating cardiac tissue to cyclic mechanical stimulation, expression of key genes associated with cardiomyocyte differentiation was evaluated. Cultured cardiac tissues under dynamic cyclic strain exhibited sustained and synchronized beating, whereas tissues maintained under static conditions showed reduced contractile coordination over time. Mechanically stimulated tissues showed increased expression of early cardiac transcription factors GATA4 (Fig. 6c) and Nkx2.5 (Fig. 6d) as compared to decellularized tissue tissue without mechanical stimulation. We found ∼50% increase expression of Nkx2.5 and ∼30% increase in the expression of GATA4 in mechanically stimulated tissues as compared to decellularized tissue without mechanical stimulation.

**Figure. 6.**
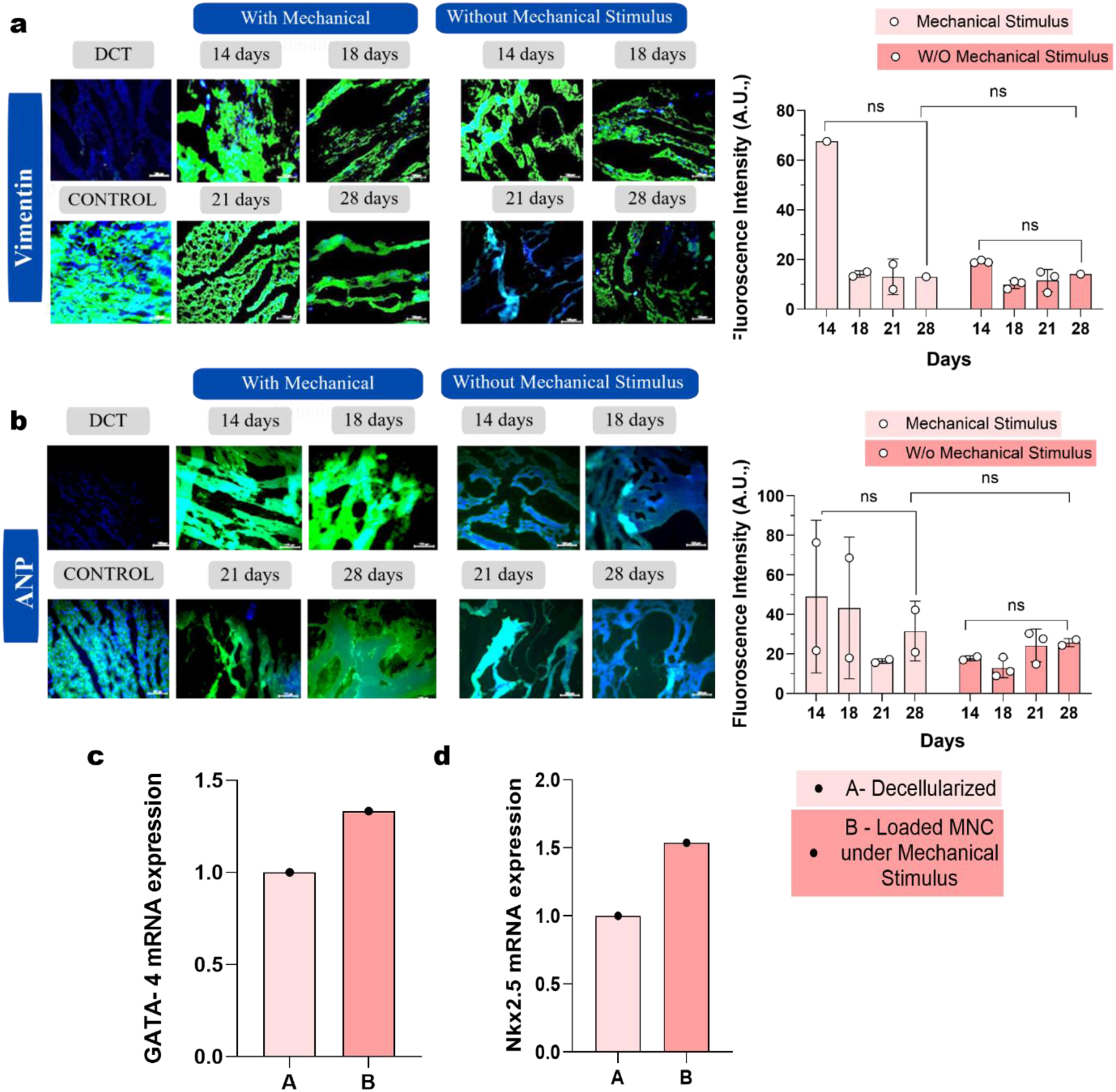
Analysis of cardiac specific-lineage under mechanical stimulation. **(a-d)** Representative immunofluorescence images and quantification of **(a)** Vimentin and **(b)** ANP expression across the culture period from day 14 to day 28. Relative mRNA expression of the early cardiac transcription factors **(c)** *GATA4* and **(d)** *Nkx2.5* in mechanically stimulated cardiac tissue compared with decellularized (acellular) scaffold controls, determined by quantitative real-time PCR (qRT-PCR). Gene expression levels were calculated using the 2^ (-ΔΔCt) method. Mechanically stimulated tissues showed a 1.5-fold and 1.3-fold increase in *Nkx2.5* and *GATA4* expression, respectively, relative to the decellularized condition. Data are presented as mean ± s.d. from [n] independent experiments.

### Influence of isoprenaline on engineered cardiac tissue beating phenotype

Isoprenaline was administered to stimulate the engineered cardiac tissue and evaluate its rhythmic beating patterns. The study aimed to characterize the dose-dependent chronotropic response and determine the concentration of isoprenaline at which arrhythmogenic activity could be induced in the engineered cardiac tissue In this representative case, FFT analysis showed that cells subjected only to mechanical stimulation exhibited a dominant low-frequency peak at approximately 0.1 Hz, whereas the addition of a low drug concentration (mention the concentration) shifted the dominant peak to approximately 0.3 Hz and produced multiple spectral components across a wider frequency range. Consistently, PSD analysis demonstrated that the mechanically stimulated condition had power concentrated primarily at low frequency, while the mechanical stimulus plus low-drug condition displayed a broader power distribution extending approximately from 0.3 to 4 Hz. This indicates that low-dose drug exposure altered the frequency composition of mechanically induced cellular motion, introducing faster and more heterogeneous temporal fluctuations. The autocorrelation profiles further showed distinct recurrence patterns: the drug-treated condition exhibited broader oscillations and a pronounced positive correlation near 7–8 s, whereas mechanical stimulation alone produced a different pattern of alternating positive and negative correlations. Thus, low-dose isoprenaline treatment modified the mechanically-induced motion signature and reduced its concentration around a single slow timescale, producing more complex and potentially less coordinated dynamics.

In another set of experiments, FFT analysis showed that mechanical stimulation alone produced a dominant low-frequency component at approximately 0.1 Hz, whereas the addition of a high drug dose (mention the concentration) generated a series of pronounced, regularly spaced peaks at approximately 0.6–0.7 Hz and its higher-frequency multiples. The corresponding PSD profile confirmed that spectral power under high-dose treatment was distributed across these repeated frequency components suggesting that the drug shifted mechanically induced cellular motion from a slow dominant timescale toward faster, strongly periodic dynamics. The autocorrelation analysis supported this interpretation: the high-dose condition showed a clear positive recurrence peak at a time lag of approximately 1.6 s, which corresponds closely to the period associated with the fundamental frequency of approximately 0.6 Hz. Beyond this recurrence, its ACF remained close to zero with a slight negative trend, indicating limited similarity over longer time intervals. In contrast, mechanical stimulation alone exhibited more irregular long-lag autocorrelation fluctuations. Collectively, high-dose of isoprenaline exposure displayed substantially modification of the mechanically-induced motion signature and introduces a faster, more pronounced periodic component

Direct comparison of the two drug doses in this representative case revealed a clear dose-dependent alteration in the temporal characteristics of mechanically induced cellular motion. FFT and PSD analyses showed that the low-dose condition contained a relatively broad and irregular distribution of frequency components, with its strongest response near approximately 0.3 Hz. In contrast, the high-dose condition displayed a series of prominent and regularly spaced peaks, beginning near 0.6–0.7 Hz and extending to higher frequencies, indicating the emergence of a faster and more strongly periodic motion pattern. The autocorrelation results supported this distinction: the high-dose condition exhibited a pronounced recurrence peak at approximately 1.6 s, consistent with the period corresponding to its fundamental frequency, followed by correlations close to zero at longer lags. Conversely, the low-dose condition retained broader and more irregular long-timescale autocorrelation fluctuations, suggesting a more complex temporal structure. Collectively, this representative comparison suggests that increasing the drug dose shifts mechanically induced cellular motion from a broadly distributed, heterogeneous frequency pattern toward a faster and more clearly defined periodic response.

**Figure. 7.**
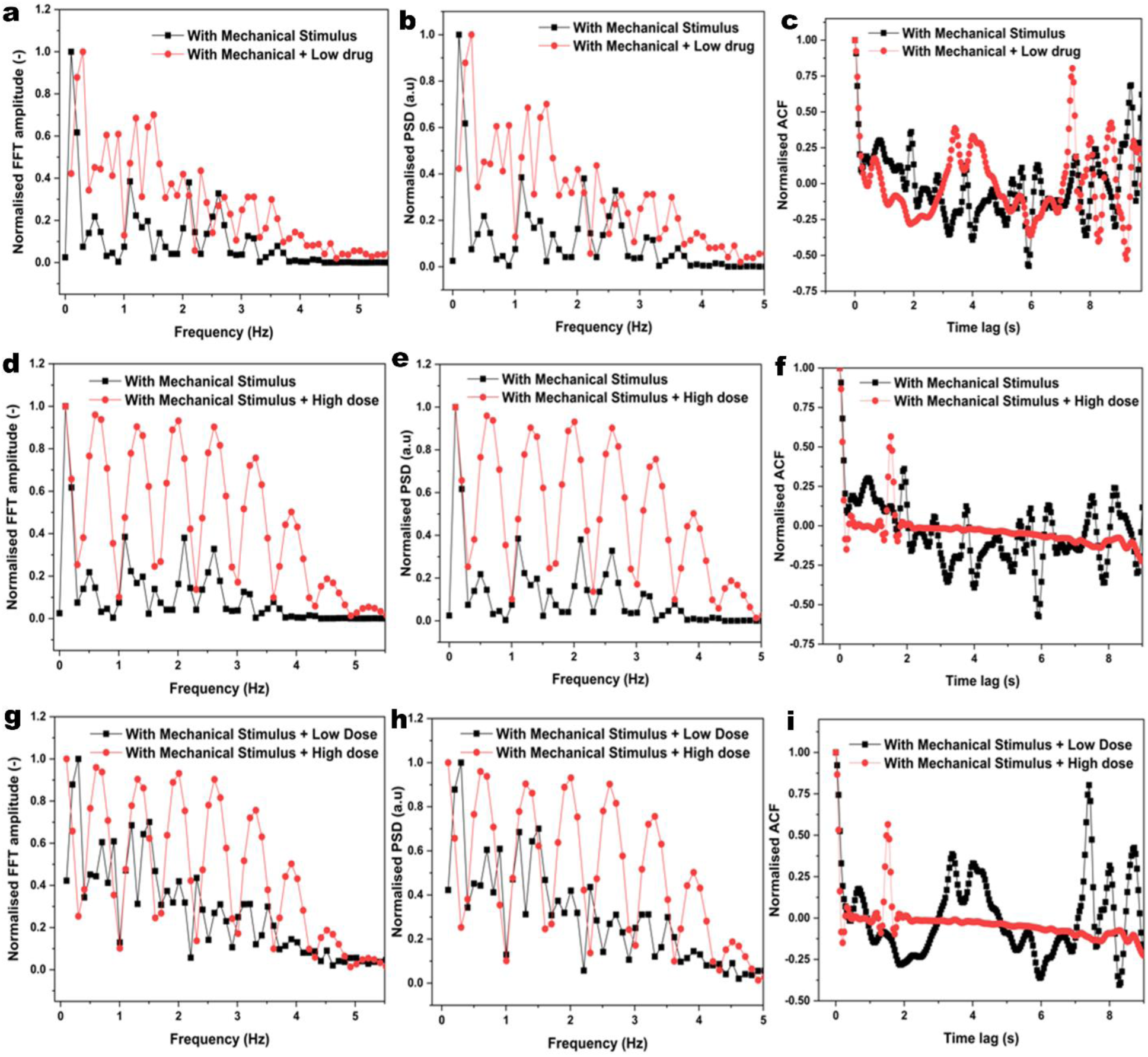
Representative frequency and time-domain analyses of cellular motion under mechanical stimulation and isoprenaline treatment. **(a–c)** Comparison between mechanical stimulation alone and mechanical stimulation with a low drug dose, **(d–f)** comparison between mechanical stimulation alone and mechanical stimulation with a high drug dose and **(g–i)** comparison between low- and high-dose drug treatments under mechanical stimulation.

## DISCUSSION

This study establishes a scaffold-level, balloon-catheter-actuated platform for delivering cyclic multiaxial mechanical (C-MAM) loading to native, decellularized tissue constructs, and demonstrates its capacity to drive cardiomyogenic differentiation of mononuclear cells seeded onto a decellularized myocardial scaffold. In contrast to the majority of existing microphysiological systems, which apply uniaxial or biaxial strain to cell-laden hydrogels^14,18^, our approach wraps intact, ECM-preserving tissue around an inflatable balloon, so that stretch is transmitted through native tissue architecture rather than an engineered gel matrix.^19^ This distinction matters because hydrogel encapsulation, while convenient for actuation and imaging, imposes a mechanically homogeneous and often isotropic environment that does not reproduce the anisotropy, fiber alignment, and through-thickness heterogeneity of real organs. By coupling a fluid–structure interaction (FSI) computational model with direct strain-sensor measurements, we show that the platform reproduces organ-specific deformation regimes from the rapid, low-amplitude cycling characteristic of cardiac and smooth-muscle tissue to the slow, high-amplitude distension characteristic of bladder filling within a single, reconfigurable device architecture^20^. The close correspondence between predicted and measured lateral-area change across four distinct loading regimes indicates that the numerical model can be used prospectively to estimate strain fields (radial, axial, circumferential) and local stresses that are not directly accessible through surface measurement, which is a practical advantage for tuning stimulation protocols to a target tissue without exhaustive empirical calibration^21,22^.

Interestingly, C-MAM conditioning of mononuclear-cell-seeded acellular myocardial scaffolds promoted markers of cardiomyocyte lineage specification and maturation increased troponin and α-sarcomeric actinin expression, a shift in Cx43^23–25^ from diffuse cytoplasmic to membrane-associated clustering, declining vimentin and ANP expression, and upregulation of the early cardiac transcription factors GATA4 and Nkx2.5, culminating in spontaneous, synchronized contractile activity that was not observed under static culture. This progression is consistent with the broader principle that mechanical loading is not merely permissive but instructive for cardiac tissue maturation, and it extends previous hydrogel-based mechano conditioning work by showing that a comparable maturation trajectory can be achieved when cells differentiate within a decellularized native matrix under multiaxial, rather than uniaxial, cyclic strain^26^. The motion-analysis data (FFT, power spectral density, and autocorrelation) further suggest that mechanical stimulation shifts cellular motion toward a slower, more coordinated low-frequency regime, and that this signature is pharmacologically tunable: isoprenaline exposure produced dose-dependent shifts toward faster, more periodic beating, providing preliminary evidence that the platform can support chronotropic drug-response assays in addition to tissue-maturation studies^27^.

The strain and stress distributions resolved by the FSI model add mechanistic detail to this picture: across all four organ regimes, the balloon–tissue interface consistently represented the region of greatest local perturbation, with elevated circumferential strain and stress progressively attenuating toward the outer tissue surface. The same interface-to-surface gradient held despite the very different amplitudes, frequencies, and thickness responses of the heart, lung, smooth-muscle, and bladder models points to a common relationship between applied pressure waveform, balloon-wall deformation, and resulting tissue-level mechanical response, one that a single validated device architecture can reproduce across mechanically distinct organ systems, with continuous perfusion supporting the nutrient and oxygen delivery needed for longer-term culture. These findings position the platform as a candidate shared tool for comparative mechanobiology, provided the same depth of validation is extended beyond cardiac proof-of-concept for cardiac tissues as reported here.

Beyond cardiac modelling, the modularity and tunability of the platform enable its application to a wide range of dynamic tissues of the body. By adjusting actuation parameters, scaffold and tissue conditioned media selection, the system can be readily adapted to study mechanotransduction, disease progression, and drug responses in organ-specific contexts. Overall, this work establishes a generalizable organ-on-chip framework that bridges native tissue mechanics with controlled microfluidic engineering, offering a powerful tool for advancing mechanobiology and translational tissue research. Holistically, this system may enable evaluation of candidate drugs without/minimal animal experiments and expedite the drug development process significantly.

## MATERIALS AND METHODS

### Design and fabrication of the device

The device was fabricated using a combination of 3D printing and soft lithography techniques (Fig. 1d). A master mold was 3D-printed using Vero thermoplastic material. To prepare the polydimethylsiloxane (PDMS) cast with a provision for catheter balloon insertion, a mixture of PDMS and curing agent in a 10:1 ratio was poured over the 3D-printed master, which was equipped with a support structure (L × W × D: 45 mm × 38 mm × 11 mm). The support structure served two key purposes: first, it provided a boundary around the master to retain the PDMS mixture in the desired shape until cured, and second, it included two hollow square features (L × W × D: 10 mm × 10 mm × 8 mm) on either side to securely position the catheter balloon. These hollow square structures were designed to accommodate tubes with diameters ranging from 2 mm to 2.5 mm.

The PDMS-curing agent mixture was poured into the master-support assembly in two stages. In the first stage, the mixture was poured and cured up to the level where the square boxes begin. This is followed by degassing and curing the assembly in a hot air oven at 70 °C for 30 minutes. In the second stage, the mixture was poured exclusively into the square boxes, followed by degassing and curing again under the same conditions for 30 minutes. Once fully cured, the PDMS cast was peeled off, tubes were pulled out of the cast and holes were punched at the inlet/outlet of the channels to enable fluid flow. The PDMS cast was then bonded to a glass slide *via* plasma treatment, and the balloon catheter was inserted into the PDMS cast. Finally, the assembly, comprising the PDMS cast, bonded glass slide, and inserted balloon catheter, was sealed with a top cap made from 3D-printed Vero Ultra Clear material (Fig. 1f).

### Numerical simulation

The designed device was numerically simulated using COMSOL Multiphysics software to model the stress-train behaviour of the myocardium tissue under cyclic mechanical stimulus. A 2D axisymmetric fluid-structure interaction (FSI) model has been considered for the purpose as shown in Fig. S3.

The problem consists of expansion/contraction behaviour of the annular air domain under positive/negative external pressure stimuli at inlet (Fig. S3). The cyclic mechanical pressure mimicking real world physiological conditions result in transverse displacement of the lateral surface which is transferred to the tissue structure wrapped around. Thus, there exists a fluid-structure interaction coupling between the air domain and the myocardium tissue as shown in Fig. S3. In present experimental study, the lateral surface of the air domain was made up of medical grade pure silicone over the decellularized tissue is wrapped. To numerically model these conditions, we consider a hyperelastic model for the lateral silicone surface while an elastic (Hookean) model was assumed for the decellularized tissue. In addition, we consider a perfect contact between the tissue and lateral surface such that the former closely mimics the motion of the latter. The material properties considered in present numerical study are adopted from previous literature and is summarized in Table S2.

To model the fluid flow behaviour in the air domain we consider continuity equation and Navior-Stokes equation given as:

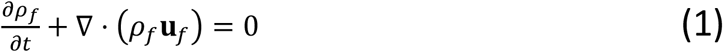

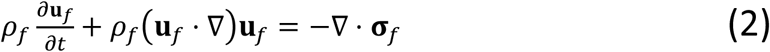

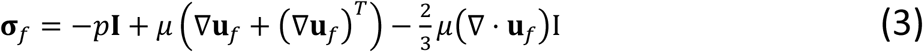

This fluid stress results in deformation of the solid objects, which is modelled using the solid mechanics equations given as:

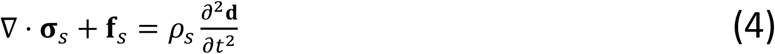

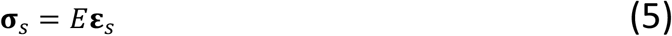

for elastic myocardium tissue

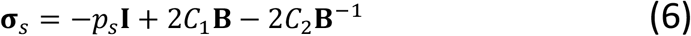

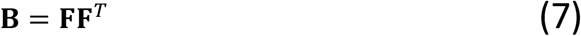

for hyperelastic silicone

The fluid and solid motions are coupled at the fluid–structure interface by enforcing continuity of velocity and stress. The corresponding fluid–structure interaction (FSI) equations at the solid– liquid interface are given as:

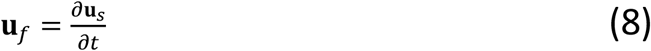

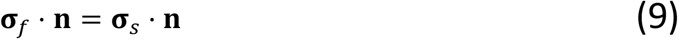

Equations (1)–(3) describe conservation of mass and momentum in the air domain and define the fluid stress tensor. Equation (4) governs solid motion, while Eqs. (5)–(7) describe the elastic tissue response, the hyperelastic silicone response, and the associated deformation tensor. Equations (8) and (9) enforce velocity compatibility and traction equilibrium at the fluid–solid interface. For the fluid domain, we consider a single inlet pressure boundary condition with all other surfaces being blocked as shown in Fig S3. The remaining fluid boundaries are treated as rigid walls, on which we apply no-slip and no-penetration condition. The interaction between the fluid and the solid is captured at the continuity of common fluid–solid interface by imposing FSI boundary conditions.

These conditions enforce continuity of velocity and displacement, along with equilibrium of traction, thereby ensuring that the hydrodynamic stresses exerted by the fluid are fully transmitted to the solid. Also, we consider the solid deformation influences the fluid motion in a fully coupled manner. For the solid domain, a fixed boundary condition is applied at the top surface, where all translational degrees of freedom are constrained to prevent rigid body motion and to provide structural support. The remaining exposed surfaces of the solid are modelled as free boundaries, on which zero external traction is applied, allowing the solid to deform naturally in response to the fluid-induced loads. Thus, these boundary conditions ensure a physically consistent representation of the coupled fluid flow and solid deformation under physiologically relevant conditions.

The coupled fluid–structure interaction problem is solved using a fully coupled manner based on Arbitrary Lagrangian–Eulerian (ALE) framework using COMSOL Multiphysics^28,29^. The fluid flow is governed by the incompressible Navier–Stokes equations, which are discretized in space using the finite element method with appropriate velocity–pressure interpolation to ensure solution stability. The solid equations are also discretized using a similar finite element framework. Finally, the fluid and solid subproblems are solved in a monolithic manner, where the governing equations of both domains are assembled into a single system and solved simultaneously at each time step. The time integration is performed using an automatic time step based on backward difference formula (BDF) with its maximum value being limited to 0.1s. The meshing operation was performed using unstructured triangular mesh with finest mesh (∼200 times finer than the tissue diameter) being located near the solid-liquid interface where FSI boundary condition is applied. To ensure numerical convergence, the tolerance value was set to 0.0005 for all variables. Also, the streamline diffusion method was selected to stabilize the solution as the numerical approximation approaches the exact solution. To verify the mesh independence conditions, we vary the number of grid points in the computation domain and an optimum mesh structure is chosen. Also, mesh deformation in the fluid domain is handled using an elastic or Laplacian smoothing approach to maintain mesh quality during large solid deformations. Convergence at each time step is ensured through nonlinear iterations until prescribed residual tolerances are satisfied.

To validate the present numerical scheme, we compare the strain at the lateral surface of myocardium tissue with the experimental results. For this purpose, a strain sensor (connected to the Multichannel Data Acquisition System) was attached to the lateral surface of decellularized tissue and the strain data was measured under mechanical stimuli similar to stretching/relaxation pattern of cardiac tissue of a healthy heart. The mechanical stimulation profiles and corresponding strain measurements acquired using the NI-9169 data acquisition system are presented in Fig. 2c, 2i, 3c, and 3i. We compare this experimental strain data with the numerical strain at lateral surface of tissue.

### Decellularization of cardiac tissue

The details of each stage of the workflow are described in Fig.1. Cardiac tissue was harvested immediately after euthanasia and transported in sodium bicarbonate-containing Ringer’s lactate solution. Following removal of excess adipose and connective tissue and rinsing with sterile distilled water, samples were sectioned (2–3 mm thickness) and incubated in antibiotic-supplemented Hank’s Balanced Salt Solution (HBSS) at 4 °C for 48 h.

Decellularization was carried out using 1% deoxycholic acid (DCA, pH 7.8)^30^ at 4 °C for 48 h, to ensure complete cellular removal. Samples were then washed thoroughly with phosphate-buffered saline (PBS) and treated with heparin (5000 U in Ringer’s lactate) for 8 h at 15–20 °C to impart anticoagulant surface properties.

Decellularized tissues were sequentially fixed in 4% and 8% formalin (pH 7.4; overnight and 8 h, respectively, at 15–20 °C), with intervening PBS washes. Samples were subsequently treated with 1% glutamic acid (pH 7.8, 1 h, room temperature). Finally, scaffolds were washed with PBS, equilibrated in DMEM before cell seeding.

### Histological Analysis

Samples were fixed in 4% paraformaldehyde, dehydrated through a graded ethanol series (70– 100%), and paraffin-embedded. To ensure representative sampling of the tissue core, samples were bisected and oriented with the inner surface facing the mold base prior to sectioning at 10 µm thickness. Sections were stained with hematoxylin and eosin (H&E) and DAPI fluorescence to assess residual nuclear/DNA content, confirming the extent of decellularization. Additional histological characterization was performed using Masson’s trichrome staining to evaluate collagen distribution and Elastic Van Gieson (EVG) staining to assess elastin fiber architecture and overall extracellular matrix integrity^31–33^.

### DNA Extraction and Quantification

Heart tissue (∼100 mg; native and decellularized) was homogenized in 0.1 M EDTA and lysed with 10% SDS, followed by centrifugation (10,000 rpm, 10 min). The supernatant was extracted with an equal volume of phenol: chloroform (1:1) and centrifuged (10,000 rpm, 10 min), then re-extracted with chloroform: isoamyl alcohol (24:1) under identical conditions. DNA was precipitated from the aqueous phase with two volumes of chilled ethanol (10,000 rpm, 10 min), washed with 75% ethanol (10,000 rpm, 10 min), air-dried, and resuspended in 50 µl TE buffer^34,35^. DNA concentration was determined and purity was assessed with a (NanoDrop Lite Plus Spectrophotometer, Thermoscientific).

### DNA Fragment Analysis

DNA fragment length was assessed by agarose gel electrophoresis. Samples (20 µl) were mixed with 4 µl of 6X loading dye and resolved on a 1% agarose gel stained with ethidium bromide (EtBr), alongside a 1 kb DNA ladder (1 Kb Plus DNA Ladder(250 μg)-10787018, Invitrogen). Gels were imaged using an E-Gel Imager (12003153-ChemiDocTM Imaging System, BioRad).

### Engineered tissue culture & initiation of mechanical Stimulus in microfluidic device

Peripheral blood mononuclear cells (PBMCs) were obtained commercially from HiMedia (India), thawed, and pre-cultured in RPMI medium (11875093-rpmi 1640 medium, Gibco) for 12–24 h prior to seeding. Approximately 2 × 10⁶ mononuclear cells were seeded into decellularized scaffolds by injection through multiple micro-punctures using 28G insulin needles, and the seeded scaffolds were maintained in complete DMEM medium (AL149A Dulbecco’s Modified Eagle Medium(DMEM),Low glucose /HiMedia) for 3 days to allow initial cell attachment^36^.

Scaffolds were then wrapped around the balloon catheter housed in the central chamber of a custom microfluidic platform to apply cyclic mechanical stimulation. A parallel set of MNC-seeded scaffolds was cultured under identical conditions but without mechanical loading, serving as a static control. Both groups were nourished using a conditioned medium containing DMEM, 10% FBS (10270106-Fetal Bovine Serum, qualified, Brazil) and 10% serum from cardiac ischemia patients. The collection of serum samples from the cardiac ischemia subjects was approved by the Institute Ethics Committee of Indian Institute of Technology Madras, Chennai (IEC/2025-03/SKD/19). The conditioned medium was administered every second day throughout the culture period, with the aim of providing pathophysiological relevant cues to support cardiomyogenic differentiation.

Mechanical stimulation was applied at 60 cycles per minute for 2 weeks. Stimulation was then withdrawn, and cultures were maintained for a further 2 weeks to allow maturation, with medium changes continuing on alternate days and periodic observations made throughout this period^37,38^. The complete experimental workflow is summarized in Fig. S2.

### Tissue harvesting, cryosectioning & immunofluorescence experiments

Cardiac tissues were carefully retrieved at 14^th^, 18 ^th^, 21^st^ and 28^th^ day and briefly rinsed in PBS, and embedded in OCT (optimal cutting temperature) compound (tissue freezing medium Brand:Tissue Tek, Cat No: (03800732)). Embedded tissues were snap-frozen in liquid nitrogen stored at −80 °C until sectioning. Cryosections (10 µm thickness) were cut using a cryostat ( Leica CM 1950) maintained at −20 °C, mounted onto glass slides, and either processed immediately for immunostaining or stored at −80 °C until further use^39^.

Sections were fixed with 4% paraformaldehyde for 15 min at room temperature, washed with Phospahte Buffered Saline, and permeabilized using 0.2% Triton X-100 for 15 min. Non-specific interactions of the antibodies were blocked by incubation of the samples with 3% bovine serum albumin for 60 min at room temperature. The tissues were then incubated overnight at 4 °C with primary antibodies specific to the structure or protein of interest (as mentioned below). Cardiomyocyte differentiation of the seeded mononuclear cells was confirmed by immunostaining for a panel of cardiac structural and functional markers. Sarcomeric organization and contractile phenotype were visualized using rabbit monoclonal anti-troponin and α-sarcomeric actinin, antibodies (1:100; PA5-28964, Invitrogen and 1:100; MA122863, Thermo Fisher, respectively) while intercellular electrical coupling was examined using rabbit monoclonal anti-connexin antibody (1:250;MA5-49139, ThermoFisher). Atrial natriuretic peptide (ANP) expression was used to assess cardiomyocyte-specific secretory phenotype using rabbit monoclonal anti-ANP antibody (1:250; PA5-72527, ThermoFisher), and Vimentin staining was included to distinguish differentiated cardiomyocytes from residual undifferentiated or mesenchymal-like cells using anti-vimentin antibody (1:250; :PA5-27231, ThermoFisher) as per the the protocol^40^.

Following primary antibody incubation, cells were washed twice with PBS and incubated overnight at 4 °C with the corresponding secondary antibodies (A-11008, ThermoFisher). Nuclei were counterstained with DAPI (ThermoFisher). Fluorescence images were acquired using an inverted fluorescence microscope equipped (Carl Zeiss Axio Vert.A1) and processed using ZEN software (Zeiss) and ImageJ (NIH).

### Analysis of gene expression in Engineered Cardiac Tissue (ECT)

RNA was extracted from engineered cardiac tissues using RNAqueous kit (Qiagen). Briefly, on day 18, tissues were added immediately to lysis buffer in microcentrifuge tube. RNA was then extracted by following the manufacturer’s instructions on the RNA aqueous kit, followed by DNAse I digestion. The yield and quality of RNA were assessed with Nanodrop spectrophotometer (Nano Drop Lite Plus Spectrophotometer – ThermoScientific). RNA isolated from ECT was reverse-transcribed to cDNA using Takara cDNA kit (RR037A). The cDNA library products were then 10-fold diluted in sterile water for quantitative PCR analysis. Expressions of the cardiac transcription factors GATA-4 and Nkx2.5^41^ were assessed by qPCR using the following primer sequences: GATA-4 forward primer (5′-GGCTATGTCCACCCCGCTCTG-3′) and reverse primer (5′-TGGCAGTTGGCACAGGAGAGG-3′) and Nkx2.5 forward primer (5′-CCCCTGGATTTTGCATTCAC-3′) and reverse primer (5′-CGTGCGCAAGAACAAACG-3′).

### Isoprenaline-induced β-adrenergic stimulation of Engineered Cardiac Tissue

Engineered cardiac tissues were treated with isoprenaline to assess β-adrenergic responsiveness^42^. Working concentrations of [0.01-0.1mg/ml /50mg of tissue] was prepared from 2mg/ml of Isoprenaline stock, which was added in the culture medium immediately after preparation to minimise oxidative degradation of the drug. Tissues were equilibrated under baseline conditions for 15 min, after which prescribed dose of Isoprenaline was administered by direct addition to the culture well. Contractile parameters (e.g., beat rate, contraction amplitude) were recorded continuously before and after drug administration, with cumulative dose-response measurements obtained by sequential addition of increasing isoprenaline concentrations, allowing [10 min] for steady-state response at each dose. Controls were run in parallel under identical culture conditions.

### Image analysis

Fluorescence images were analyzed using ImageJ software (National Institutes of Health, USA)^43^. For quantification, raw images were imported without prior contrast adjustment, and thresholding was applied uniformly across all images within a given staining set to distinguish positive signal from background. Nuclear counterstaining (DAPI) was used to determine total cell number, allowing marker-positive cells to be expressed as a percentage of total nuclei. Quantification was performed on a minimum of three independent fields per sample, averaged across biological replicates.

### Data and statistical analysis

Direct comparisons were made using unpaired Student’s t-test. When comparisons were made across more than two groups, one-way analysis of variance (ANOVA) was first performed to determine global significance before performing post-hoc tests for pairwise differences with the Holm-Sidak method, unless noted otherwise in specific figure captions. All curve fitting was done using GraphPad Prism. Gene expression data were statistically analysed with GraphPad Prism. Significance was determined using P < 0.05.

## Acknowledgements

This work was financially supported by The Indian Council of Medical Research, Government of India (Grant number: IIRPIG-2023-0000716). We also gratefully acknowledge the financial support from ICMR and IIT Madras. We thank Prof. Dhiman Chaterjee and Prof. Tuhin Subra Santra for their valuable suggestions in developing the design of microfluidic device. We also thank Sabari (Research Intern) and Shubham Kumar Mishra (Research Scholar) and for their valuable contributions to image quality enhancement.

