## Supplementary information for "An Efficient Microfluidic Platform for Organ-Specific Cyclic Multiaxial Loading and Differentiation of Progenitor Cells"

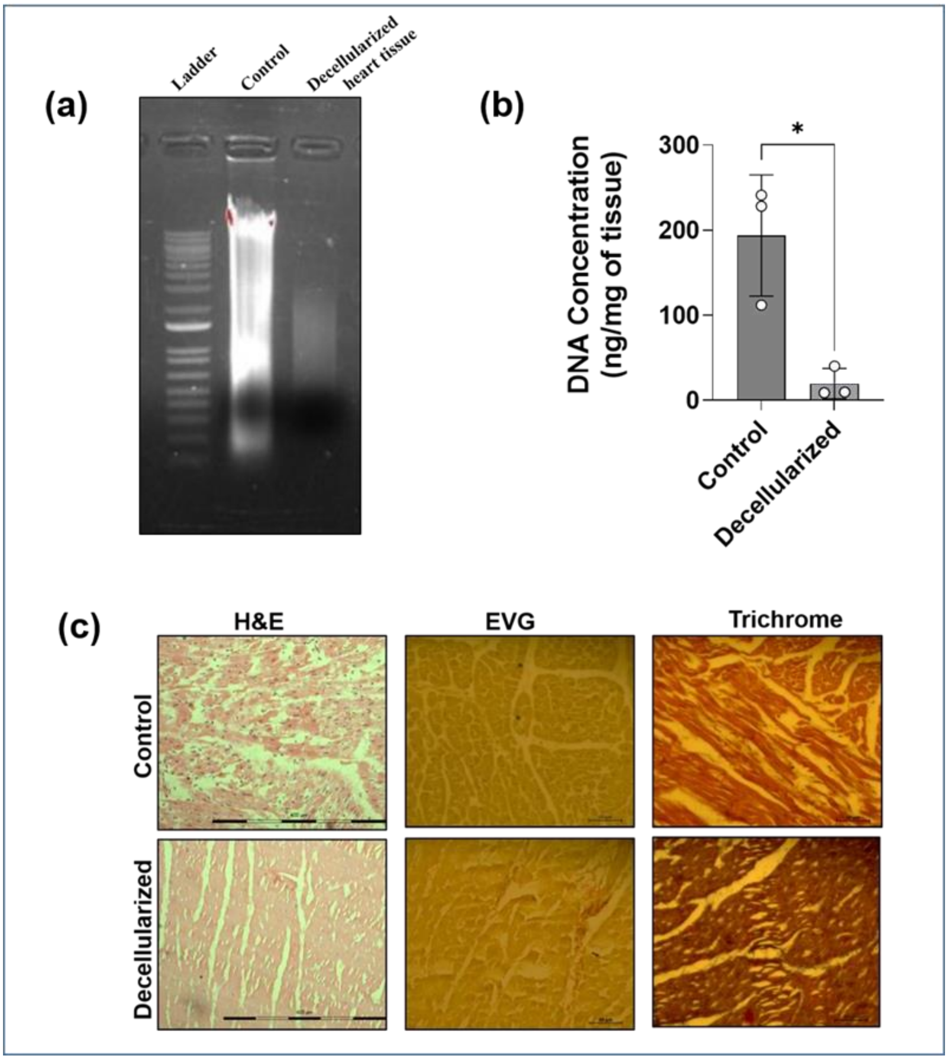


**Figure. S1 | Validation of decellularization efficiency by gel electrophoresis and immunohistology. (a)** Agarose gel electrophoresis (0.8%) showing removal and fragmentation of DNA in decellularized tissues compared with native controls, and (ii) quantitative measurement of residual genomic DNA content in native and decellularized cardiac tissues. (d) Histological evaluation of decellularized cardiac tissue sections stained with hematoxylin and eosin (H&E), Elastic Van Gieson (EVG), and Masson's trichrome, confirming effective cellular removal with preservation of extracellular matrix architecture, including collagen and elastic fibers. Data are presented as mean ± s.d. Statistical significance was assessed by t-test (control vs. decellularized) (*$p < 0.05$).

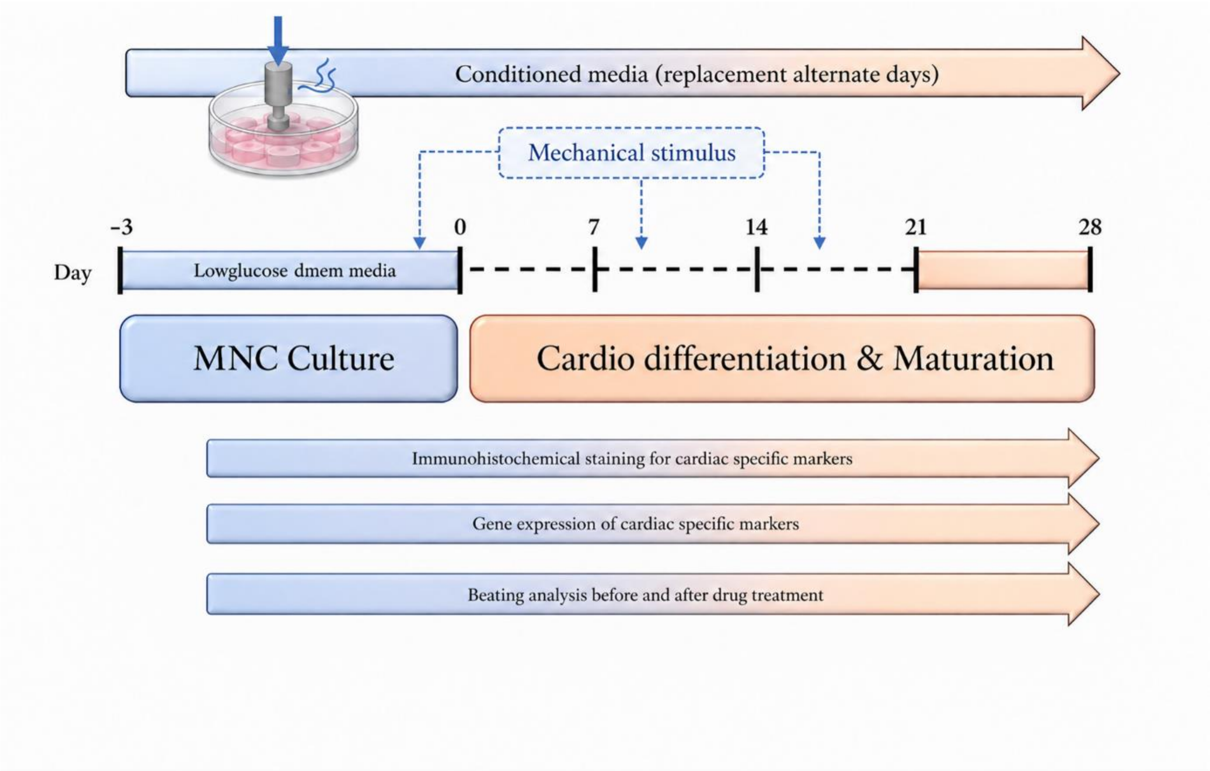


**Figure. S2 | Schematic workflow of cell culture and differentiation.** The schematic summarizes the experimental sequence from cell seeding through differentiation, maturation and downstream analysis.

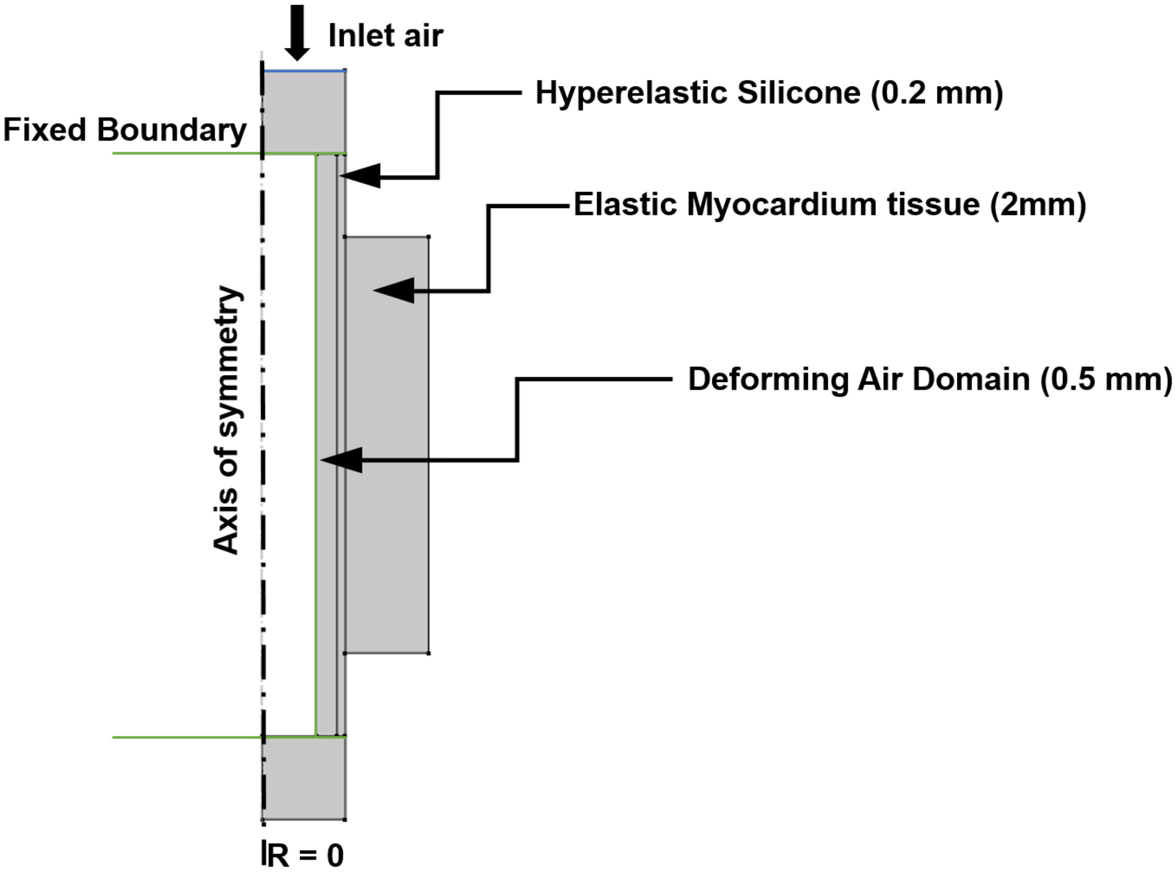


**Figure. S3 |**. Axisymmetric model of the air–silicone–myocardium system, showing layer dimensions and boundary conditions.

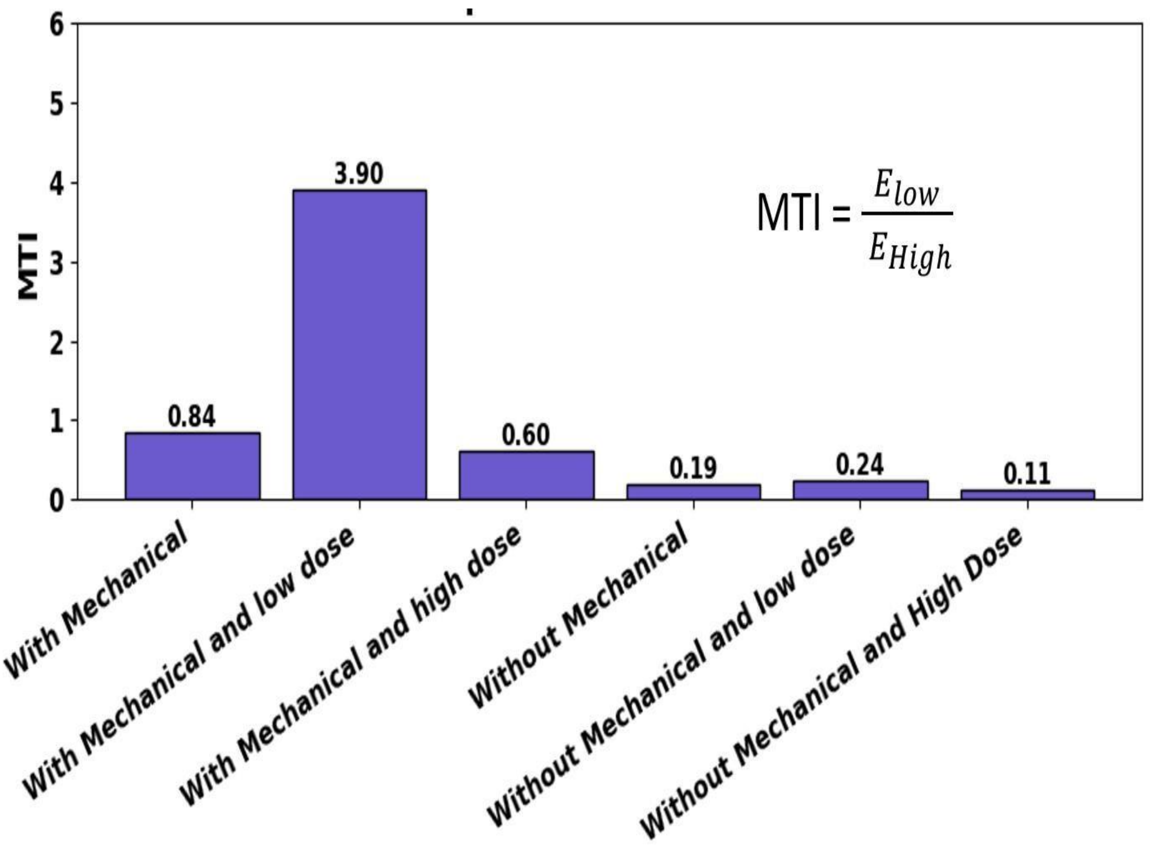


**Figure. S4 | Mechanotransduction Index (MTI =** $E_{low}/E_{high}$**)** for all experimental conditions, comparing mechanical stimulation and drug dose variations.

The Mechanotransduction Index (MTI), defined as the ratio of low-frequency to high-frequency spectral energy ($E_{\square\square\square}/E_{h\square\square h}$), is employed to quantify the degree of coordinated contractile behavior across all experimental conditions. As illustrated in Fig. S4, the highest MTI value is observed for the condition combining mechanical stimulation with a low drug dose (MTI ≈ 3.90), indicating a strong dominance of low-frequency components and, consequently, highly coordinated and organized contraction. Mechanical stimulation alone results in a moderate MTI (≈ 0.84), suggesting that mechanical cues enhance coordination but are insufficient to achieve optimal functional organization in the absence of biochemical support. In contrast, increasing the drug dose under mechanical stimulation leads to a reduction in MTI (≈ 0.60), implying that excessive biochemical influence disrupts coordinated behavior.

Conditions without mechanical stimulation consistently exhibit low MTI values (≈ 0.11–0.24), irrespective of drug dose, reflecting a predominance of high-frequency fluctuations associated with uncoordinated and noisy activity. Notably, the lowest MTI is observed for the combination of no mechanical stimulation and high drug dose (≈ 0.11), indicating severely impaired contractile coordination. Overall, these findings demonstrate that mechanical stimulation plays a primary role in enhancing contractile organization, while an optimal (low) drug dose further amplifies this effect. In contrast, high drug concentrations and the absence of mechanical cues lead to diminished coordination and increased irregularity, underscoring the importance of balanced mechanochemical coupling in regulating cardiac-like contractile behavior. The Mechanotransduction Index (MTI), defined here as the ratio of low- to high-frequency spectral energy, is conceptually related to LF/HF-type spectral indices used in cardiovascular variability analysis to distinguish organized low-frequency oscillations from high-frequency noise. In agreement with reports that dynamic mechanical loading enhances maturation and synchrony in engineered cardiac tissues, while cardioactive drugs exert dose-dependent effects on beat-rate variability in stem-cell-derived cardiomyocytes, our data show that mechanical stimulation markedly increases MTI and that an optimal (low) drug dose further amplifies this effect, whereas high drug concentrations and the absence of load are associated with reduced MTI and poorly coordinated contractions[1-3].

Overall, the results indicate that mechanical stimulation serves as a primary driver of structural and contractile coordination, while an appropriate (low) drug dose further enhances this effect, promoting optimal functional maturation. In contrast, elevated drug concentrations and the absence of mechanical cues result in reduced coordination and increased irregularity, highlighting the essential role of balanced mechanochemical coupling in governing cardiac-like contractile behavior.

**Table S1. Heart: Stress distribution through the tissue thickness at 0.2 s**

| **Tissue location** | **Radial stress, kPa** | **Axial stress, kPa** | **Circumferential stress, kPa** |
|---|---|---|---|
| Tissue–balloon interface | -13.6 | 76.4 | 185 |
| Mid-thickness | -0.113 | -0.055 | 0.380 |
| Outer tissue surface | -0.010 | -0.047 | 0.260 |

**Table S2. Lungs: Stress distribution through the tissue thickness at 0.6 s**

| Tissue location | Radial stress, kPa | Axial stress, kPa | Circumferential stress, kPa |
|---|---|---|---|
| Tissue–balloon interface | -14.817 | 93.550 | 226.480 |
| Mid-thickness | -0.145 | 0.069 | 0.484 |
| Outer tissue surface | -0.0129 | 0.059 | 0.3248 |

**Table S3. Smooth Muscle: Stress distribution through the tissue thickness at 1.05 s**

| Tissue location | Radial stress, kPa | Axial stress, kPa | Circumferential stress, kPa |
|---|---|---|---|
| Tissue–balloon interface | -12.299 | 72.914 | 175.190 |
| Mid-thickness | -0.1064 | 0.052 | 0.3591 |
| Outer tissue surface | -0.00839 | 0.0444 | 0.2453 |

**Table S4. Bladder: Stress distribution through the tissue thickness at 20 s**

| Tissue location | Radial stress, kPa | Axial stress, kPa | Circumferential stress, kPa |
|---|---|---|---|
| Tissue–balloon interface | -14.829 | 93.578 | 226.560 |
| Mid-thickness | -0.145 | 0.069 | 0.481 |
| Outer tissue surface | -0.0137 | 0.0601 | 0.326 |
